# Investigations into modifications of neuromuscular physiology by axonal transport disruptions in *Drosophila* SOD1 mutants

**DOI:** 10.1101/2020.11.27.399956

**Authors:** Tristan C. D. G. O’Harrow, Atsushi Ueda, Xiaomin Xing, Chun-Fang Wu

## Abstract

The reactive oxygen species (ROS)-scavenging enzyme Cu/Zn superoxide dismutase (SOD1) is an evolutionarily conserved mechanism for the maintenance of oxidative homeostasis, and missense mutations in the human SOD1 gene are associated with the motor neuron degenerative disease amyotrophic lateral sclerosis (ALS). Mutations in the *Drosophila melanogaster* SOD1 gene (*Sod*) shorten fly lifespan, attenuate motor function, and induce developmental mortality. We have previously found morphological and physiological defects at the neuromuscular junctions (NMJs) of hypomorphic *Sod*^*n108*^ and ALS model *Sod*^*G85R*^ mutant larvae. Here, we report genetic interactions causing striking modifications of *Sod* mutant phenotypes by mutations in the gene *Prickle* (*Pk*), which are linked to planar cell polarity, epilepsy, and axonal transport disruptions. *Pk* is expressed as two isoforms *prickle-spiny-legs* (*sple*) and *prickle-prickle* (*pk*), which are each suppressed by specific hypomorphic mutations (*sple1* and *pk1*). Interestingly, *Sod* phenotypes are distinctly modified depending on both the *Pk* isoform suppressed, and whether said suppression is heterozygous or homozygous. Heterozygous *sple1* and *pk1* improved the developmental survival of *Sod* mutants, whereas homozygous *sple1* and *pk1* drastically increased mortality. Further, only heterozygous *sple1* and *pk1* clearly ameliorated morphological defects at the neuromuscular junctions of *Sod* mutant larvae. Pharmacological treatment of *Sod* mutants reveals allele-specific motor neuron terminal hyperexcitability, characterized by synaptic transmissions of extended duration and abnormal presynaptic Ca^2+^ transients. Heterozygous *sple1* mutation suppresses this *Sod* mutant terminal hyperexcitability, but *pk1* does not. We reversed this suppressive effect of *sple1* by pharmacological blockade of Ca^2+^-activated K^+^ channel *slowpoke*. Altogether, this study builds on our prior knowledge of *Sod* mutant development and physiology to show that axonal transport-linked gene mutations strikingly modify *Sod* mutant phenotypes, providing strong evidence for a role of intracellular transport in alterations of neuromuscular morphology and physiology by SOD1 mutations.

## INTRODUCTION

The ROS-scavenging enzyme Cu/Zn superoxide dismutase (SOD1) is highly conserved across phyla, and is expressed in a wide variety of tissues (Uhlén et al., 2015). Consequently, mutations in the gene encoding SOD1 have pervasive consequences for physiology, developmental viability, and organismal longevity (Phillips et al., 1989).

Research continues to reveal defects in motor neuron development and physiology associated with mutations in the SOD1 gene, including hypomorphic mutations and mutations linked to familial amyotrophic lateral sclerosis (FALS) (Rosen et al., 1993).

Reactive oxygen species (ROS) are known to behave as signaling molecules to play a wide variety of roles in healthy neuronal development and physiology (Thannickal & Fanburg, 2000; Milton & Sweeney, 2011). During neurodevelopment, ROS modify the biochemistry of cytoskeletal components to influence their dynamics and structure (Wilson & González-Billaut, 2015; Munnamalai & Suter, 2009). The reactive superoxide anion (O_2_^-^) has also been shown to regulate synaptic plasticity (Klann et al., 1998; Massaad & Klann, 2011). In *Drosophila* larval motor neurons, oxidative stress influences synaptic structure and transmission (Sanya et al., 2002), and regulates activity-dependent neuronal plasticity (Oswald et al., 2018).

Loss of SOD1 function due to genetic mutation allows for oxidative stress in excess of physiological levels. Resulting physiological disruptions include the misfolding of proteins (Berlett & Stadtman, 1997), peroxidation of lipids (Altan et al. 2003), and alteration of nucleic acids (Ali et al., 1996; Aitken & Krausz, 2001). High oxidative stress affects neurodevelopment by disrupting cytoskeletal structures (Landino et al., 2002; Munnamalai & Suter, 2009). SOD1 has been shown to support axon outgrowth and survival, and oxidative stress induced by loss of SOD1 or treatment with a redox toxin triggers neurodegeneration. (Fischer & Glass, 2010). Further, SOD1 knockout mice display NMJ neurotransmission impairments and neuromuscular degeneration (Shi et al., 2014). The *n108* allele (Campbell et al., 1985) of the *Drosophila* gene for SOD1 (*Sod*) is functionally null, and *n108* mutant larvae and flies display hypersensitivity to oxidative stress (Phillips et al., 1989). The adults also exhibit severely reduced lifespans (Phillips et al., 1989) and motor defects (Ruan & Wu, 2008). We have also previously shown that *n108* mutant larvae display altered synaptic transmission at the neuromuscular junction, and supernumerary axon discharges (Ueda et al., *manuscript in preparation*), and defective neurotransmission in the adult giant fiber escape circuit (Ueda et al., 2021).

Through mechanisms that remain to be clarified, SOD1 is linked to FALS when specific amino acid substitutions are present. Mouse, *C. elegans*, and *Drosophila* models expressing ALS-linked mutant SOD1 exhibit motor dysfunction or lethal paralysis (Wang et al., 2008; Wang et al., 2009; Watson et al., 2008; Sahin et al., 2017; Agudelo et al., 2020). Prior to detectable neurodegeneration, ALS mutant SOD1 impacts neurodevelopment and cell metabolism. ALS mutant SOD1 localizes to actin structures, altering growth cone dynamics and eventual neuronal morphology (Osking et al., 2019). Misfolded mutant SOD1 protein also co-localizes with mitochondria in the axons of murine motor neuron, where mitochondrial morphology is altered (Vande Velde et al., 2011) and mitochondrial Ca^2+^ buffering capacity is defective (Damiano et al., 2006). The *Drosophila* ALS-linked *Sod* mutant *G85R* displays defective larval locomotion, and consistently fails to eclose from the pupal stage (Held et al., 2019; Sahin et al., 2017). *G85R* larvae also display proprioceptor neuron defects and associated motor circuit dysfunction (Held et al., 2019).

In a separate report, we have characterized *Sod* mutant allele-dependent defects in NMJ physiology and synaptic morphology (Ueda et al., Manuscript in preparation). In screening for modifiers of synaptic transmission in *Drosophila Sod* mutants, we fortuitously found genetic interactions between *Sod* and the planar cell polarity (PCP) and axonal transport-linked gene *prickle* (*pk*) (Gubb & García-Bellido, 1982; Gray et al., 2011; Lin & Gubb, 2009; Tree et al., 2002). *pk* is expressed in *Drosophila* in the form of the Pk^Pk^ (Prickle Prickle) and Pk^Sple^ (Prickle Spiny-legs) isoforms. Isoform-specific loss-of-function mutants (*pk1* and *sple1*, Gubb et al., 1999) display distinctly altered microtubule-mediated transport and microtubule dynamics in larval motor axons, with related changes in neuronal excitability (Ehaideb et al., 2014; Ehaideb et al., 2016). We have previously found that *pk1* and *sple1* mutant larvae exhibit alterations in motor neuron development and synaptic transmission that may be downstream of axon transport modifications (Ueda et al., 2022).

Here, we report the findings that *pk* isoform loss-of-function (LOF) mutations either improve or reduce the developmental viability of *Sod* mutant *Drosophila* pupae in a genetic dosage-dependent manner. Similarly, only heterozygous *pk* LOF suppresses NMJ morphology defects in *Sod* mutants. We also demonstrate that *pk* LOF induces allele-specific alterations *Sod* mutant NMJ transmission. Additionally, we show that selective pharmacological blockade of ion channels reveals NMJ terminal hyperexcitability in *Sod*^*G85R*^, and that this phenotype is suppressed by partial LOF of the Pk^Sple^ isoform of *pk*. This work therefore demonstrates that dysregulations of development and neuromuscular physiology induced by *Sod* mutation can be altered by *pk* isoform dose manipulation, indicating potential roles of modified intracellular transport in the generation of *Sod* mutant phenotypes.

## MATERIALS & METHODS

### Fly stocks

Flies and larvae were housed in bottles containing a cornmeal-agar medium (Frankel & Brousseau, 1968), supplemented with 3g of yeast paste prepared as a 1:1 mixture of ddH_2_O and active dry yeast (Red Star Yeast Co., LLC). Flies were reared at a room temperature of 22 ± 1°C. The *Sod* mutants examined were those of alleles *n108* (Phillips et al., 1989) (gift from John Phillips, University of Guelph, Ontario, Canada) and *G85R* (şahin et al. 2017) (gift from Barry Ganetzky, University of Wisconsin-Madison, USA). The wild-type control strain used for Figures 1, 2, 3, and 4 was Canton-S (CS). For experiments in which a motor neuron membrane tag was needed, the stock used carried a chr. II *UAS-CD8-GFP* recombined with a chr. II motor neuron GAL4 driver (Brand & Perrimon, 1993), either *C164-Gal4* (Torroja et al., 1999) or *OK371-Gal4* (Mahr and Aberle, 2006). The original recombined *OK371-Gal4 UAS-CD8-GFP* (*OK371::CD8-GFP*) was obtained from the National Centre for Biological Sciences (Bangalore, Karnataka, India). *Sod* lines used to produce the data represented in Figure 1 were balanced over a TM6 balancer with a dominant *Tubby* (*Tb*) marker (TM6, *Tb*). The *Pk*^*Pk*^ and *Pk*^*Sple*^ lines used in this study have been documented previously (Ehaideb et al., 2014). The wild-type control strain used for Figure 5 was the Oregon-R (OR) used in Ehaideb et al. (2014). All *Pk;Sod* double-mutant lines were generated in the Wu lab by conventional genetic crosses.

**Figure 1.**
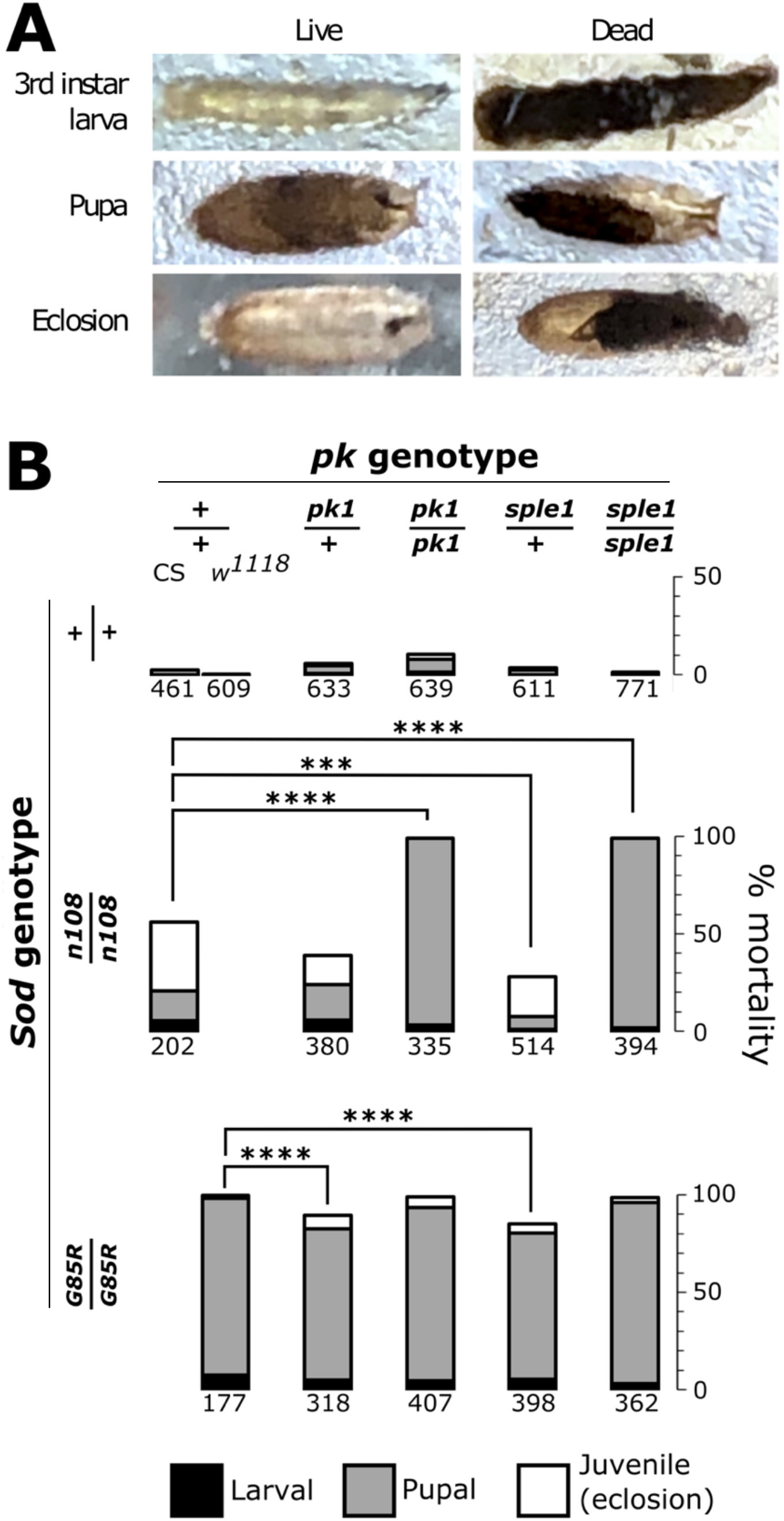
Developmental lethality in *Sod* mutants is modified by *pk* mutations. (A) Representative images of live and dead larvae and pupae, an empty pupal casing left after successful eclosion of the juvenile fly, and the carcass of a juvenile dead during eclosion. (B) *Sod* mutant survival during development is improved by one copy of either *sple1* or *pk1*. Bars show percentages of individuals dead at the 3rd instar larval stage (black), during pupal development (gray), and at eclosion from the pupal casing (white). Genotypes shown are *Sod* mutants *G85R* and *n108, pk* mutants *pk1* and *sple1, pk1/+* and *sple1/+* heterozygotes, and double-mutants of those *Sod* and *pk* alleles. Both wild-type CS and *w1118* are included as controls due to differing genetic backgrounds of *G85R* and *n108. n* indicated below each bar. \*\*\**P* < 0.001, \*\*\*\**P* < 0.0001, Fisher’s exact test. Statistics are for cumulative developmental mortality.

**Figure 2.**
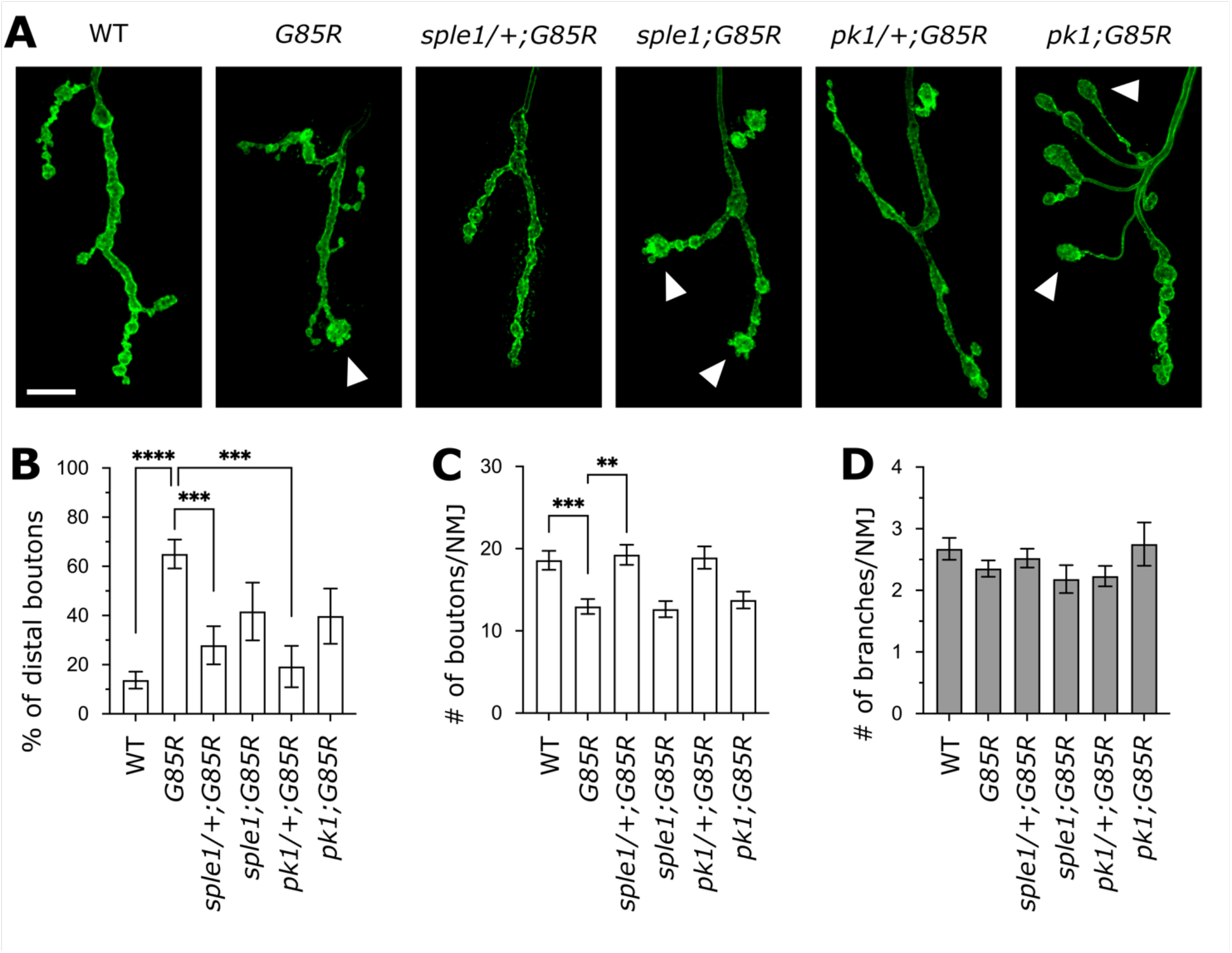
Heterozygous *prickle* loss-of-function suppresses *Sod*^*G85R*^ neuromuscular junction morphology defects. (A) Representative images of type Ib NMJs from muscle 4 in wild-type (WT), *Sod*^*G85R*^ (*G85R*), heterozygous and homozygous double-mutants of *sple1* with *G85R* (*sple1/+;G85R* and *sple1;G85R*), and heterozygous and homozygous double-mutants of *pk1* with *G85R* (*pk1/+;G85R* and *pk1;G85R*). Immunostaining shows presynaptic morphology (Anti-HRP). White arrows indicate *Sod* phenotypic enlarged distal boutons. Scale bar = 10 μm. (B) Percentage of distal synaptic boutons that are enlarged in each genotype (see methods). (C) Numbers of presynaptic boutons (see methods). (D) Numbers of presynaptic branches (see methods). Bars in B, C and D indicate mean, error bars indicates SEM. *N* of NMJ = WT: 46, *G85R*: 34, *sple1/+;G85R*: 23, *sple1;G85R*: 11, *pk1/+;G85R*: 13, *pk1;G85R*: 12. *\*\*P* < 0.01, *\*\*\*P* < 0.001, *\*\*\*\*P* < 0.0001, one-way ANOVA followed by Tukey’s test.

**Figure 3.**
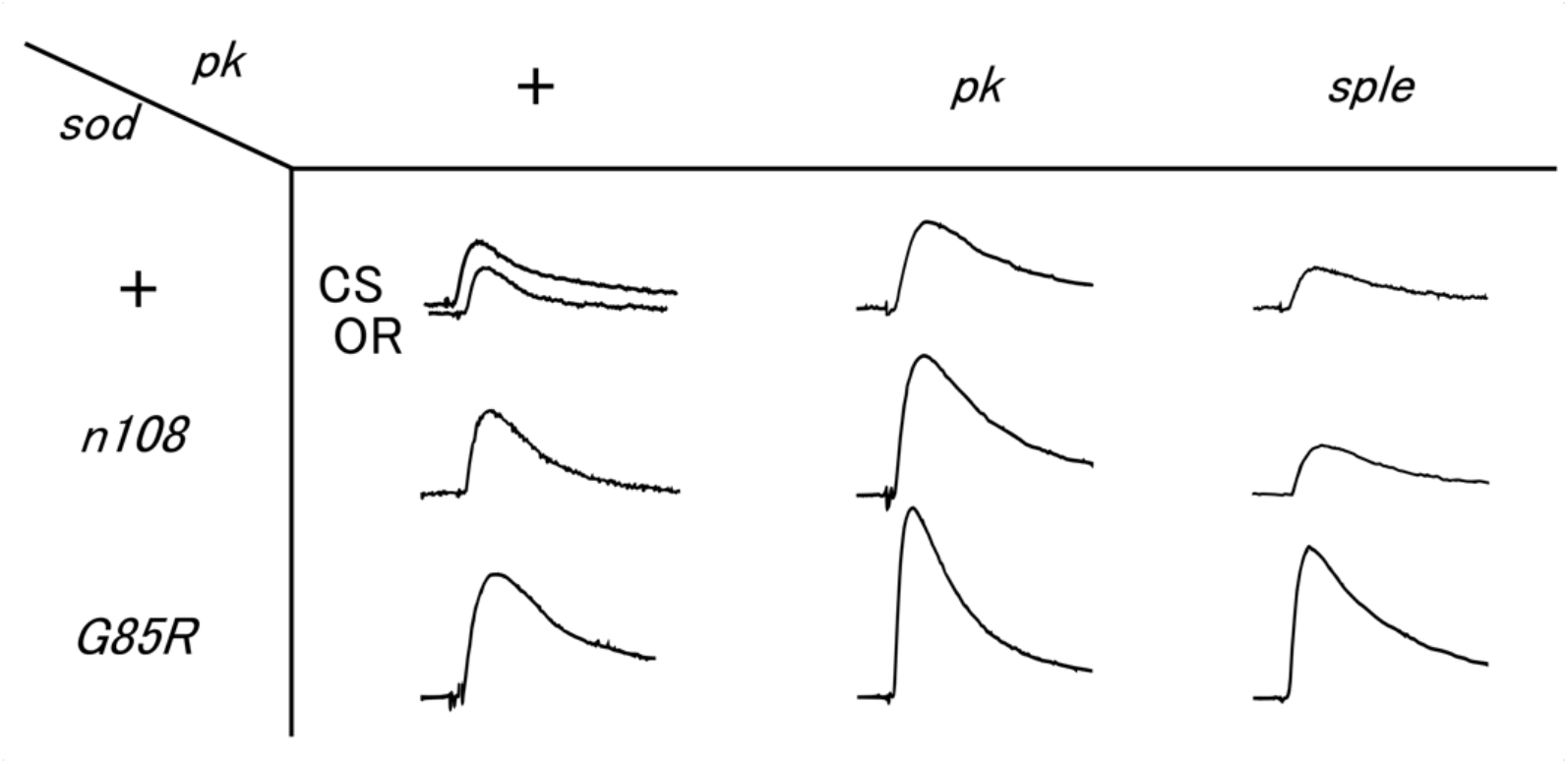
Homozygous *pk* mutations alter the size of excitatory junctional potentials (EJPs) from body wall muscles in *Sod* mutant larvae. Two different WT controls (CS and OR) are shown. EJPs were measured in HL3.1 saline containing 0.2 mM Ca^2+^.

**Figure 4.**
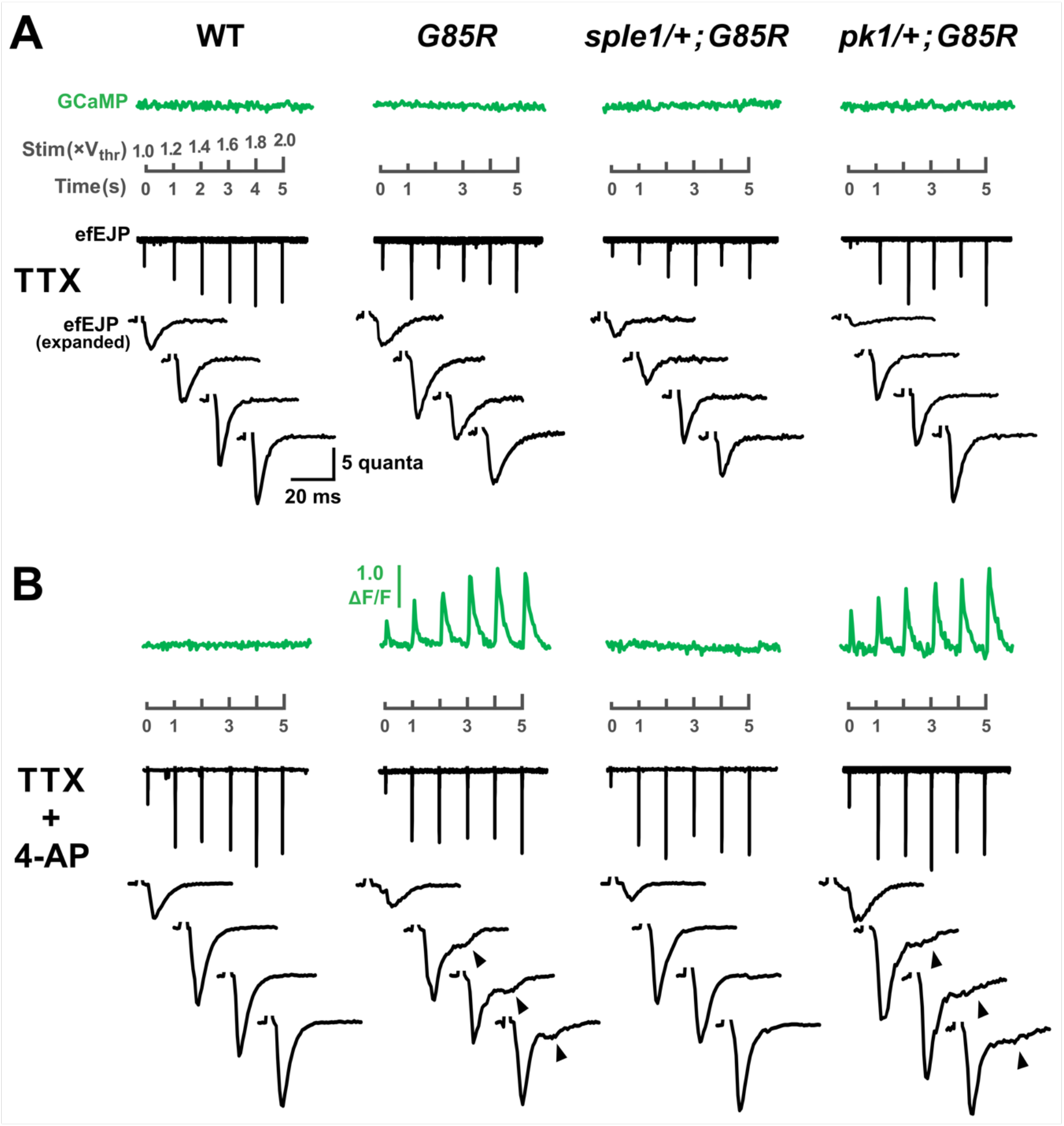
Heterozygous *sple1* suppresses the NMJ terminal hyperexcitability of *Sod*^*G85R*^. (A) efEJP of *sple1/+;G85R* and *pk1/+;G85R* (with TTX alone, see methods) are qualitatively similar to *G85R* and WT. Stimuli 1 ms in duration were delivered at 1 Hz, increasing in intensity from 1× to 2× threshold voltage. GCaMP (top, green) fluorescence remains at baseline during stimulation in all genotypes. Gray scale bar between the GCaMP and efEJP traces indicates time scale, time of stimulus (1/second), and stimulus intensity for the GCaMP and top efEJP trace. Expanded efEJP traces are responses at 0, 1, 3, and 5 s. (B) Presynaptic GCaMP spikes and efEJP extension are absent in *sple1/+*;*G85R*, indicating that one copy of the *sple1* mutant allele suppresses *Sh* blockade-induced terminal hyperexcitability of *G85R*. Retention of 4-AP-induced *G85R* hyperexcitability characteristics (GCaMP spikes and extended efEJP) in *pk1/+*;*G85R* demonstrates that the interaction of *sple1* with *G85R* is *prickle* allele-dependent. Arrowheads indicate efEJP ‘shoulder’ characteristic in *G85R* and *pk1*/+;*G85R*.

**Figure 5.**
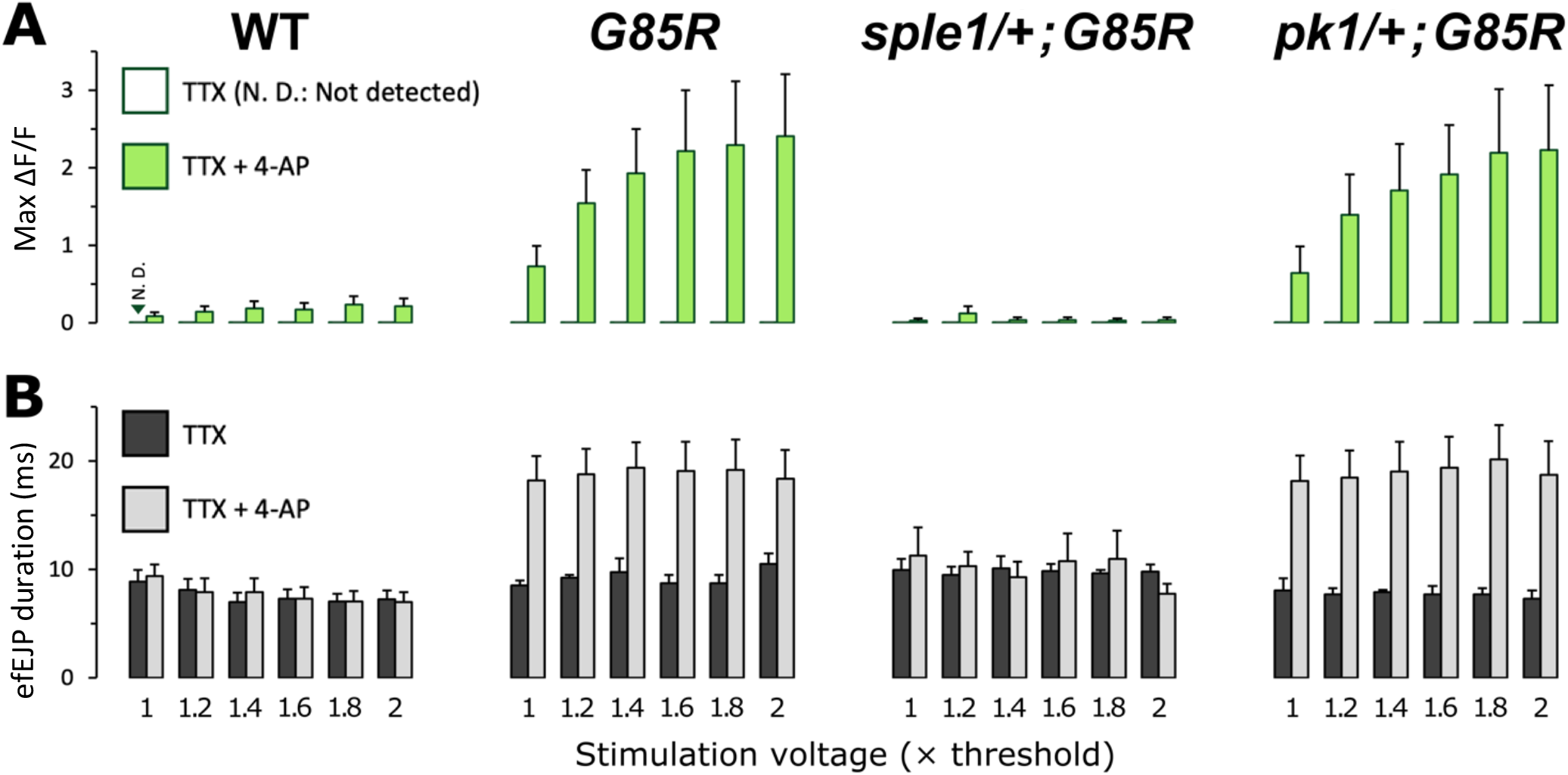
Quantification of presynaptic GCaMP signal amplitude and efEJP duration in *G85R* and *pk*/+;*G85R* double-mutants. (A) Mean GCaMP signal amplitudes (peak ΔF/F) in response to 1 Hz stimuli, of intensity from 1× to 2× threshold voltage. Error bars indicate SEM. In the presence of TTX alone, GCaMP signals were always undetectable (no deviation from baseline). Thin dark green line indicates baseline. 4-AP induced large single stimulus-evoked GCaMP spikes (light green bars) in *G85R* and *pk1*/+;*G85R* larvae. WT and *sple1*/+;*G85R* larvae typically lacked these spikes. See GCaMP traces in Figure 6. (B) Mean durations (ms) of the efEJP co-occurring with the GCaMP signals quantified above. efEJP traces were measured at 20% amplitude (see methods). WT and *sple1*/+;*G85R* larvae displayed efEJP of similar duration in the absence (dark gray bars) or presence (light gray bars) of 4-AP. *G85R* and *pk1*/+;*G85R* efEJP under 4-AP were typically about twice the duration of efEJP without 4-AP. The efEJPs of extended duration were those displaying the shoulder-like characteristic indicated in Figure 6.

### Developmental lethality studies

Housing and medium conditions for the developmental lethality study (Figure 1) were identical to those described in the main ‘Fly stocks’ section. Flies including 20-40 females were used to seed each bottle. Counts of deaths and eclosions were performed 16-17 d. after seeding. The TM6, *Tb* balancer carried by *Sod* heterozygous wandering 3rd instar larvae and pupae was used to distinguish them from *Sod* mutant homozygotes. Dead larvae appear as darkened corpses on the wall of the housing bottle. Dead pupae were entirely black, or contained a mixture of dark and empty patches, as confirmed by transmitted light under a dissection scope. Juvenile flies dead during eclosion were distinguished from flies in the process of eclosing by a second observation 15-30 min after the first. Juvenile fly corpses were also frequently darker than their live counterparts.

### Larval preparations, vital staining, & immunohistochemistry

Wandering third instar larvae were dissected, as per the ‘fillet preparation method’, in HL 3.1 saline (Feng et al., 2004). HL3.1 used in immunohistochemistry or vital staining experiments contained 0.1 mM Ca^2+^. Tetramethylrhodamine (TMRM, C_25_H_25_ClN_2_O_7_) (Floryk & Houštěk, 1999) vital staining of larval preps for mitochondria was done at 100 nM in HL 3.1 for 20 minutes (in a light-impermeable chamber to prevent TMRM photobleaching), after which the bath was replaced with TMRM-free HL 3.1 before microscopy. Before antibody staining, larval preps were fixed in 3.7% formaldehyde in HL 3.1 for 25-30 minutes. Fixed preps were incubated with an FITC-conjugated goat antibody to HRP at 1:50 in phosphate buffer saline, at 4°C for 12-48 h. Anti-HRP (Jan & Jan, 1982) recognizes a neural carbohydrate antigen in *Drosophila* (Kurosaka et al., 1991). For double-staining with anti-HRP and anti-Dlg (monoclonal AB 4F3: Parnas et al., 2001) or anti-Futsch (monoclonal AB 22C10: Fujita et al., 1982; Zipursky et al., 1984) staining, fixed larval preps were incubated with the primary monoclonal AB at 1:100 in phosphate buffer saline, at 4°C for 12-48 h, rinsed, and incubated in phosphate buffer saline containing anti-HRP(conj. FITC) and a secondary TRITC-conjugated anti-mouse IgG antibody, both at 1:50. The 4F3 and 22C10 antibodies were obtained from the Developmental Studies Hybridoma Bank (DSHB) (University of Iowa, Iowa City, IA, USA). The FITC-conjugated anti-HRP and TRITC-conjugated anti-mouse IgG were obtained from Jackson ImmunoResearch Laboratories (West Grove, PA, USA). Larval NMJ images, vital and fixed, were gathered on a Leica DM IL LED inverted microscope (Leica Microsystems Inc., Buffalo Grove, IL, USA), using the Leica Application Suite X software.

### Larval NMJ morphology and mitochondrial distributions

Examination of *Sod* single-mutant NMJ morphology (Figure 2) was carried out in muscles 4 (M4) and 13 (M13), in abdominal segments A3, A4, and A5, and quantifications from the different segments were pooled. Examination of *Pk;Sod* double-mutant NMJ morphology (Figure 4) was carried out in muscle 4 (M4) in abdominal segments A2, A4, and A6, and quantifications from the different segments were pooled. Counts of boutons from M4 include only type Ib boutons. Ib and Is boutons and branches in M13 were quantified separately. Counts of specific bouton morphologies (‘unsegregated’ boutons and large terminal boutons) are from M4 (Ib). For branch counting, a branch was defined as a terminal process carrying at least two boutons (Zhong et al., 1992). A bouton was described as incompletely segregated (‘unsegregated’) if the width of the narrowest point on the neck preceding it was greater than 80% of the width of the widest point on the bouton. If multiple unsegregated boutons appeared in a row, it could be difficult count them. In these cases, counts were conservative. A terminal bouton was determined to be the ‘largest’ if it was measurably larger than any other bouton on that branch, past the last branching point.

### Electrophysiology

All electrophysiological recordings were based on published protocols, and performed in HL3.1 saline (Feng et al., 2004) (Composition (mM): 70 NaCl, 5 KCl, 0.1 CaCl_2_, 4 MgCl_2_, 10 NaHCO_3_, 5 trehalose, 115 sucrose, 5 HEPES) (pH adjusted to 7.1). Intracellular recording of excitatory junction potentials (EJPs) was adapted from Ueda and Wu (2006). Recording of extracellular focal EJPs (efEJPs) was shown in Ueda and Wu (2009) and Xing and Wu (2018a,b). For nerve-evoked action potentials, 0.1 ms stimuli were applied to the cut end of the segmental nerve through a suction pipette. To enable analysis of presynaptic terminal excitability, we also performed electrotonic stimulation (Wu et al., 1978; Ganetzky & Wu, 1982; 1983). Briefly: Tetrodotoxin (TTX) was applied to block Na+ channels. To achieve direct electrotonic stimulation of the terminal, a 1 ms stimulus was applied to the cut segmental nerve, which was sucked into the stimulation electrode, nearly up to the entry point into the muscle, so as to allow the stimulus to passively propagate to the terminal and depolarize the endplate membrane (Lee et al., 2014). In this manner, CaV channels were directly triggered by local depolarization, independent from invasion of axonal Na+ action potentials. Further, a significant portion of multiple K+ channel species, including *Shaker* (Kv1), were blocked by co-application of 4-aminopyridine (4-AP) and tetraethylammonium (TEA) (Jan & Jan, 1977; Singh & Wu, 1999). Electrotonic stimulation of the larval NMJ with TTX, 4-AP, and TEA has been shown in Lee et al. (2014). Such manipulations allow for the examination of terminal excitability driven by Ca2+ channels with reduced repolarization regulation by a range of K+ channels, with the exclusion of axonal potential interplay.

### Statistics

Total mortalities per genotype (Figure 1) were compared using Fisher’s exact test. Counts of NMJ boutons and branches and of enlarged terminal and unsegregated boutons (Figures 2 and 3) were examined with Kruskal-Wallis testing between all groups within muscle # and parameter (eg. M4, boutons), followed by rank-sum (Mann-Whitney U) post hoc between individual groups. EJP sizes were compared with one-way ANOVA (Figure 4B). Plateau-like EJP durations were compared by F-test (Figure 4D). Counts of NMJ mitochondrial macropunctae (Figure 6) were compared with Kruskal-Wallis testing followed by rank-sum (Mann-Whitney U) post hoc.

**Figure 6.**
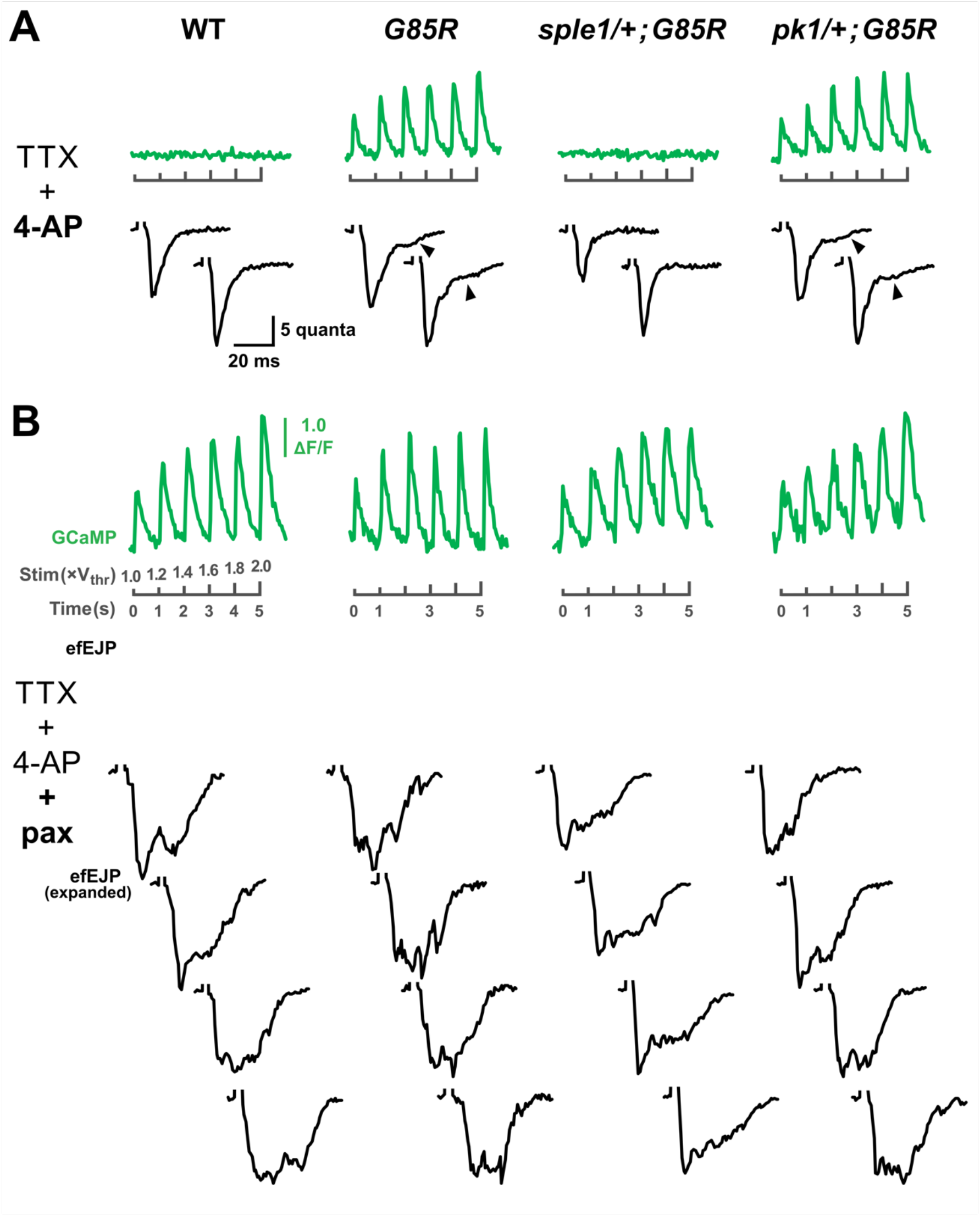
Blockade of *slowpoke* BK channels in addition to *Shaker* channels produces extreme motor terminal hyperexcitability. (A) GCaMP and efEJP traces from the TTX + 4-AP condition for WT, *G85R, sple1*/+;*G85R*, and *pk1*/+;*G85R* are shown for comparison to panel B. Expanded efEJP traces are responses at 1, 3 s. (See Figure 6 for additional and more thorough representation.) (B) Blockade of *Sh* and *slo* K^+^ channels, by 4-AP and paxilline respectively, produces extreme terminal hyperexcitability that shares characteristics with the hyperexcitability observed in *G85R* and *pk1*/+;*G85R* under the influence of 4-AP alone. Paxilline-induced terminal hyperexcitability is qualitatively similar across all genotypes examined.

**Figure 7.**
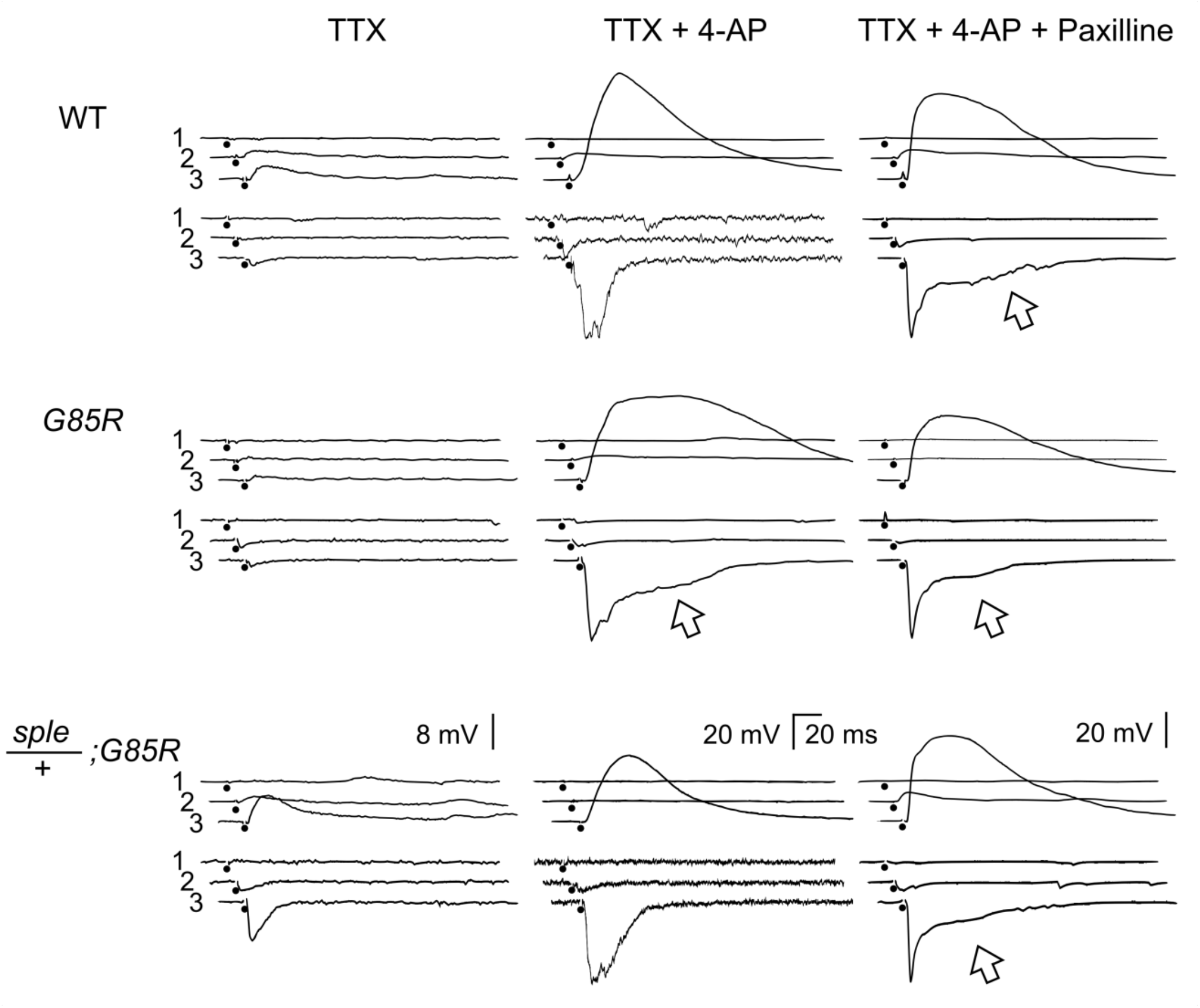
Extended NMJ transmission in *G85R* broadens whole-cell EJPs in the postsynaptic muscle, and heterozygous *sple1* ameliorates this phenotype. Double-recording of whole-cell EJPs and extracellular macropatch EJPs induced by electrotonic depolarization (•) of nerve terminals in the presence of TTX. Application of 4-AP increased the size of EJPs in all genotype. Note that *sod*^*G85R*^ exhibited broadened whole-cell EJPs associated with a shoulder of macropatch EJPs (arrow), while WT and *pk*^*sple*^/+;*sod*^*G85R*^ lacked them. Application of paxilline induced broadened whole-cell EJPs plus a shoulder of macropatch EJPs (arrow) in all genotypes. 1, 2, 3 indicates sub-threshold, near threshold, and supra-threshold (x3 ∼ x5) stimulation. Macropatch EJP traces are normalized.

## RESULTS

### Lethality of *Sod* mutants during different developmental stages

Lifespan defects in adult *Drosophila* homozygous for hypomorphic *Sod* allele *n108* (Ruan & Wu 2008) and pupal lethality in homozygotes of *n108* and the ALS-linked point-mutant allele *G85R* (şahin et al., 2017) have been previously characterized. We further noted that *n108* and *G85R* mutants exhibit lethality in different phases of the pupal stage (Figure 1B). While the majority of *n108* death occurred at eclosion from the pupal stage (Figure 1A, lower-right image), over 90% of *G85R* individuals died during pupal development (Figure 1A, middle-right image), unable to develop to the eclosion stage. Both *Sod* mutants also displayed mild levels larval lethality (Figure 1A, upper-right image), similar between the alleles. By comparison, *Drosophila* of control strains CS and *w1118* exhibited only very small levels of developmental lethality that were greatly exceeded by the significant lethality in the *Sod* mutants (p < 0.001). Heterozygous *Sod* mutants (*n108*/+ and *G85R*/+) did not differ in lethality from the control strains (Supplemental Table 1), indicating a lack of phenotypic dominance of *Sod* mutant lethality.

We found that the developmental lethality of *n108* and *G85R* mutants was modified by mutations in *prickle*. Two copies of either the *pk1* or the *sple1* allele conferred full developmental lethality to *n108* mutants, but unlike the lethality profile of the *n108* single-mutant that generally died at eclosion, the *pk1*;*n108* and *sple1*;*n108* double-mutants died during pupal development. *pk1*;*G85R* and *sple1*;*G85R* double-mutants remained as fully lethal as the *G85R* single mutant, although rate of progression to eclosion was marginally increased. Crucially, one copy of either *pk* mutant allele had the opposite effect: *Sod* mutant lethality was suppressed. One copy of *pk1* (*pk1*/+;*n108*) lowered the overall rate of *n108* lethality below 50%. A copy of *sple1* (*sple1*/+;*n108*) was even more effective, with lethality reduced below 30%. Single copies of *pk1* and *sple1* (*pk1*/+;*G85R* and *sple1*/+;*G85R*) conferred to *G85R* mutants rates of survival to adulthood of approx. 10% and 14% respectively. These small proportions are notable when compared to the complete pre-eclosion lethality of the *G85R* single mutant.

### Altered larval neuromuscular junction morphology

The *Sod* mutants displayed mild alterations in NMJ outgrowth, that were readily visible in counts of boutons per NMJ (Figure 2B). Counts of Ib branches and boutons in M4 and M13 in *G85R* yielded distributions that were not significantly different from WT. While the increase in *G85R* M13 Is branches was nonsignificant, the numbers of Is boutons were significantly increased compared to WT (p < 0.05). NMJ outgrowth was increased in *n108* compared to WT, with more more Ib boutons in M4 (p < 0.05) and M13 (p < 0.001), more Ib branches on M13 (p < 0.01), and a visible but nonsignificant increase in numbers of M4 Ib branches (Figure 2B). Where *n108* NMJ complexity was significantly greater than WT, it was also significantly increased compared to *G85R*, indicating physiological differences between mutants with the loss-of-function allele and those with the ALS-linked allele.

Both *Sod* mutants displayed increases in numbers of unsegregated boutons per M4 Ib branch compared to WT (*G85R*: p < 0.001, *n108*: p < 0.05) (Figure 2C). Similarly, terminal boutons that were enlarged compared to other boutons on the same M4 Ib branch appeared more frequently in both *Sod* mutants than in WT (*G85R*: p < 0.001, *n108*: p < 0.01) (Figure 2C).

Unlike the *Sod* single-mutants, phenotypes of NMJ complexity (M4, Ib) in *Pk*^*Pk*^, *Pk*^*Sple*^, and their double mutants with *G85R* were most visible in changes to the numbers of branches per NMJ. Compared to WT, *Pk*^*Pk*^;*G85R* displayed significantly more boutons (p < 0.01), and *Pk*^*Sple*^;*G85R* displayed significantly fewer (p < 0.01). Both double-mutants displayed fewer branches than either corresponding *Pk* single-mutant (both p < 0.001), and *Pk*^*Sple*^;*G85R* displayed a number of boutons similar to the *G85R* single-mutant, strikingly fewer than the *Pk*^*Sple*^ single-mutant. (p < 0.001). Both *Pk* single-mutants displayed slightly more boutons than WT (both p < 0.05). None of the *Pk* single-mutants or *Pk*;*G85R* double-mutants displayed a number of unsegregated boutons that was significantly different from WT, nor was there a clear difference between WT and the *Pk* single-mutants in the rate at which terminal boutons were enlarged. However, both *Pk*;*G85R* double-mutants displayed enlarged terminal boutons at rates lower than *G85R*, but still greater than WT (both p < 0.01).

### Genetic interaction between *Sod* and *Pk* in motor neuron excitability and synaptic transmission

In light of the NMJ morphological defects manifested in *Sod* through its interaction with *Pk* mutations, we examined neuromuscular transmission for related defects. When the motor axon bundle was stimulated with 0.1 ms pulses in HL3.1 saline containing 0.2 mM Ca2+, we detected reliable EJP responses in *Sod* and *Pk* mutant larvae. In *Sod* mutants (hypomorphic alleles *n108* and *x39*/*n108*, as well as the ALS-related *G85R* allele), EJP response upon nerve stimulation was similar to that in WT (p > 0.05, Figure 4B). EJP size in the *Pk* alleles of *Pk* (*Pk1* and *Pk30*, data combined in Figure 4) was comparable to WT, while EJP size was significantly decreased in the mutants of the *Sple* allele (p<0.05, One-Way ANOVA).

However, the double-mutant *Pk*^*Pk1*^;*Sod*^*G85R*^ showed unusually large EJPs (Figure 4B, p < 0.05, One-Way ANOVA). Abnormal synaptic transmission can be attributed to alterations in ion channels participating in regulation of presynaptic terminal membrane depolarization to control Ca^2+^ influx that triggers release of synaptic vesicles from presynaptic terminals.

### Sod^G85R^ allele-specific motor terminal hyperexcitability

*Sod* mutant axonal excitability having been studied (Figure 3) (Ueda et al., In preparation), we examined the isolated electrophysiological properties of the *Sod* mutant larval NMJ using a previously established protocol of direct electrotonic stimulation of the axon terminal, after silencing the axon Na+ spikes via TTX (Ganetzky & Wu, 1982, 1983; Ueda & Wu 2009; Lee et al., 2013). Varying intensities of direct electrotonic stimulation of the presynaptic terminal revealed no clear qualitative differences between *G85R, n108*, and WT in loose-patch focal recording (see methods) of efEJP (extracellular focal excitatory junction potentials) from single synaptic boutons (Figure 4A). Exposure of the larval preparation to the *Shaker* K_V_ channel blocker 4-aminopyridine (4-AP, 200 nM) induced NMJ hyperexcitability in *G85R* larvae that was typically absent under the same condition in *n108* and WT. The *Shaker* blockade-induced hyperexcitability of *G85R* manifested in the form of a shoulder-like extension of the efEJP (Figure 4B). We quantified this characteristic with a measurement of the duration at 20% amplitude beyond the baseline (Figure 6). A presynaptically expressed Ca^2+^ indicator (GCaMP6f) also communicated the state of Ca^2+^ currents in the recorded synaptic boutons during evoked neurotransmission, which revealed that *G85R* also exhibited brief presynaptic Ca^2+^ currents in response to electrotonic stimuli in the presence of 4-AP. In fact, we found that every extended efEJP was accompanied by a spike in presynaptic GCaMP (Figure 4B, Figure 6). Given this consistent association between augmented presynaptic Ca^2+^ currents and modified postsynaptic response, we suspect that the prolonged efEJP is elicited by prolonged neurotransmitter release from the presynaptic terminal.

### pk^sple1^ suppression of Sod^G85R^ hyperexcitability

The *sple1* null allele of *prickle* exerted a striking effect on the NMJ physiology of *G85R*: One copy of *sple1* was sufficient to suppress both the efEJP extension and the simultaneous presynaptic Ca^2+^ current amplification (Figure 5B, Figure 6). That the control *sple1*/+ mutant larvae exhibit mild hyperexcitability in response to 4-AP reveals that the genetic interaction between *G85R* and *sple1* is not simply the sum of opposing phenotypes. Further, the suppressive interaction of *sple1/*+ with *G85R* physiology is *pk* allele-dependent, and does not arise from the combination of the *pk1* allele with *G85R* (Figure 5B, Figure 6).

## DISCUSSION

Our study contrasts the loss-of-function *n108* allele of *Sod* with the ALS-linked *G85R* allele, by demonstrating clear differences in the developmental stages at which the mutants of each allele die. The contrast is further enhanced by differences between their neuromuscular synapse morphologies. We found potential mechanistic links between abnormal distribution of mitochondria in *Sod*^*G85R*^ and synaptic bouton enlargement (Ueda et al., Manuscript in preparation), consistent with the striking interaction with *Pk*, known to affect axonal transport. Further, following pharmacological blockade of Na+ and K+ currents in our electrophysiological experiments, the remaining Ca2+ currents appeared to initiate more physiological actions in *Sod*^*G85R*^, in a manner suppressed by *sple1* mutation.

### Developmental death in *Sod* mutants

This study of *Drosophila Sod* mutant lethality extends specificity and aspects of mutational effects on the developmental stages in which deaths occur. Specifically, we distinguish between death of the fully developed juvenile fly during eclosion from the pupal casing, and death mid-pupal development, which is characterized by a pupal casing that remains closed. As a result, we established allele-dependent developmental lethality phenotypes. The *n108* lethality occurs largely during eclosion and may be due to physical weakness of the juvenile. The vast majority of *G85R* death occurs during pupal development, often prior to formation of adult morphology, as seen by a lack of red adult eyes visible through the pupal casing of late-stage pupae (Bainbridge & Bownes, 1981).

### Aberrant NMJ morphology and physiology

Overgrowth of the *Sod*^*n108*^ NMJ has been documented previously, and another oxidative stress-sensitive mutant (*spinster*) was shown to have similar overgrowth. Further, the two genes show synergistic interactions (Milton et al., 2011). The *G85R* larval NMJ has also been characterized (Held et al., 2019) although the bouton number did not seem to differ significantly from their control. In this study, we also found no clear difference in bouton numbers, but striking bouton morphological alterations could be demonstrated in *G85R* larvae (Figure 2A). Additional immunostaining of proteins constituting the presynaptic machinery, such as synaptotagmin and dynamin (Estes et al., 1996), or postsynaptic elements such as glutamate receptors (Harris & Littleton, 2015), could elucidate the functional relevance of the unusual bouton morphologies in *Sod* mutants.

Mutant SOD1 is the most commonly identifiable risk factor in human patients with familial ALS (Zarei et al., 2015). By exploiting the amenability of the *Drosophila* larval neuromuscular preparation to the study of motor synapses at an early developmental stage, we detect clear signs of synaptic physiology and morphology alterations in *Sod* mutants.

Additionally, abnormal distributions of synaptic mitochondria in *Sod*, and severe NMJ phenotypes in *Sod* double-mutants with transport-affecting *Pk* mutations, suggest dysfunctional transport may contribute to early SOD1-linked pathology. Our work therefore adds to our understanding of phenotypes and mechanisms that will facilitate further studies to uncover critical events leading to irreversible and terminal motor neuron degeneration in ALS.

---

**Table 1.**
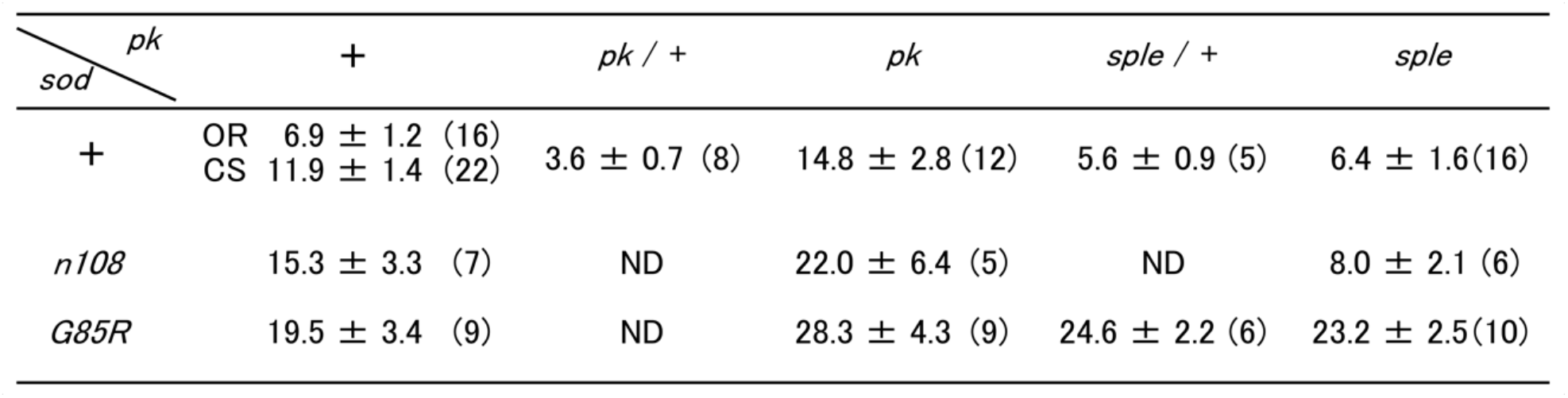
Muscle EJP size (mean ± SEM) in *Sod, pk*, and double mutant combinations. Wild-type CS is the control strain for *Sod* and wild-type OR is the control for *pk. N* of NMJs are indicated in parentheses. EJPs were measured in HL3.1 saline containing 0.2 mM Ca^2+^.

## Notes

### Competing Interest Statement

The authors have declared no competing interest.

### Summary of Updates

New figures 1-7 and addition of a table to include new data since original version submitted on 12/31/2021. The text has been updated accordingly.

